# Factors associated with adherence to the Integrated Management of Childhood Illness (IMCI) guidelines for under-five years’ old care in Burkina Faso primary health care facilities

**DOI:** 10.1101/510099

**Authors:** Sabere Anselme Traoré, Serge M.A. Somda, Joël Arthur Kiendrébéogo, Jean-Louis Kouldiati, Paul Jacob Robyn, Hervé Hien, Nicolas Méda

## Abstract

**Objective:** To assess the adherence to Integrated Management of Childhood Illness (IMCI) guidelines in primary health care facilities in Burkina Faso and to determine the factors associated.

**Materials and Methods:** We used data from a large survey on health facilities, held from October 2013 to April 2014. Primary health facilities were evaluated, health workers interviewed and consultations observed. The standard guideline for an under five year’s old child consultation was the Integrated Management of Childhood Illness (IMCI).

**Results:** 1,571 consultations were observed, carried out by 522 different practitioners. The danger signs were usually not checked (13.9% only checking for at least three general danger signs). The adherence for cough (74.8%), diarrhoea (64.9%), fever (83.8%) and anaemia (70.3%) was higher. The principal factors found to be associated with poorer adherence to guidelines of consultation were female sex (Rate Ratio (RR) = 0.91; 95% CI 0.86 – 0.95), non-nurse practitioner (RR=0.93; 95% CI 0.88 – 0.97), IMCI training (RR=1.06; 95% CI 1.01 – 1.11), non-satisfaction of the salary (RR=0.95 95% CI 0.91 – 0.99).

**Conclusion:** This study highlights a poor adherence to the IMCI guidelines and by then, revealing a poor quality of under-five care. Indeed, many characteristics of health workers including gender, type of profession, training satisfaction with salary were found to be associated with this adherence. Therefore, more initiatives aiming at improving the quality of care should be developed and implemented for improving the child health care.

## Introduction

In 1990, the mortality of under-five years’ old was really high in the Low and Middle Income Countries (LMICs). In Burkina Faso in particular, the mortality rate was 202 per 1000 living births [1]. To reverse the trend and meet the Millennium Development Goals (to reduce mortality among children under 5 years of age by two thirds over 2000), the Integrated Management of Childhood Illness (IMCI) approach was proposed in 1999 by the World Health Organization (WHO), the United Nations International Children’s Emergency Fund (UNICEF) and other technical partners. Because of its innovative implementation, the IMCI guidelines became the key strategy of child survival in most LMICs [2]. The IMCI strategy focuses on the children’s main causes of death and includes three components: (i) improving case management practices of health workers (especially in outpatient health facilities); (ii) strengthening health systems, in particular for drug supplies; and (iii) promoting community and family health practices [2–5].

A global evaluation of the IMCI in terms of impact, cost and effectiveness was conducted in many LMICs and confirmed the relevance of the strategy [6–11]. It revealed that, well implemented, the IMCI might reduce the mortality of under-five and improve their nutritional condition. Also, the investment in the strategy was profitable; as IMCI is six times cheaper than the traditional approach if health care is properly provided. Besides, it improved health workers’ performance and quality of care.

In Burkina Faso (BF), the Ministry of Health, through the Office of Mother and Child’s Health (*Direction de la Santé de la Mère et de l’Enfant*), adopted the strategy in 1999 and implemented it since 2003 [12,13]. However, in 2015, mortality among under five years old children remained high (89 per 1000 living births). Even if undeniable headways have been made (the mortality rate was reduced by 2.27 in 25 years), new Sustainable Development Goals (SDGs) targets for child mortality to end preventable deaths in children under 5 were set (reduction in neonatal mortality to as low as 12 deaths per 1,000 live births and under-5 mortality to at least as low as 25 deaths per 1,000 live births) and all the efforts to cover them must be done.

The main causes of avoidable deaths of children are well known: 27% are due to infections, 23% to malaria, 14% to acute respiratory diseases and 10% to diarrhoea, with malnutrition present in almost half of cases [1]. This suggests that there is still room for improvement in the implementation of the guidelines by identifying barriers and strengthening implementation to sustain current success.

Around the world, the IMCI guidelines’ impact on changes in care management and behaviours by health care providers are mixed in practice due to numerous barriers. While the application of the IMCI protocol to assess a patient can be completed in 10-15 minutes if health services are well organized, many health workers see patients for only a few minutes [14]. Other barriers to following IMCI guidelines include: workload, lack of motivation, difficult working conditions (unsuitable premises, sub-equipment, mismanagement of inputs and drug shortages, etc.), cumbersomeness of the IMICI training processes (lack of opportunities and staffing for training, lack of supervision) [13,15–19]. The same barriers have been observed in BF as well in many primary health care (PHC)) facilities and heavily negatively impacts the quality of care [15,20,21]. As quality of care determines demand and ultimately children’s health and survival [22–25], our study aims to assess the adherence to the IMICI guidelines by the PHC providers in Burkina Faso and to determine the factors associated with.

## 2 Materials and Methods

### 2.1 Study context and design

This study used secondary data, collected as part of the baseline survey of a performance-based financing (PBF) impact evaluation (Data in S1 Text). The pilot project was started in 2011with three health districts and then, extended to 12 other health districts in 2013. It was implemented by Burkina Faso Ministry of Health (MoH), with the technical and financial support of the World Bank through the Health Results Innovation Trust Fund (HRITF). The impact evaluation was led by the University of Heidelberg (Germany), the University of Montreal (Canada) and Centre MURAZ (Burkina Faso) [26].

It was a large scale facility-based survey, we used in this paper data regarding the working conditions and the practices of health workers (in relation to the consultations they provided to under-five years old children with reference to IMCI recommendations.

### 2.2 Study population and sampling

To conduct the impact evaluation [26], 24 health districts (out of 63) have been selected (12 intervention and 12 control districts). The 12 intervention districts (districts where the PBF strategy would be implemented) have been chosen by the MoH on the basis of poor quality outcomes for the following indicators: contraceptive prevalence, assisted deliveries, antenatal consultations and post-natal consultations. The 12 control districts (districts where the PBF strategy would not be implemented but would serve as comparison districts with the intervention ones) were chosen by the research team in charge of the impact evaluation, identified due to their relative proximity and similarity to the intervention districts. In each intervention district, all PHC facilities were included, except the newly opened ones (less than 6 months), the ones not yet opened or still under construction, or health facilities offering only certain types of services (e.g. only maternity). In each control district, PHC facilities were selected by a simple randomized draw as follow: one control health facility to four intervention health facilities.

PHC facilities (either from an intervention or control district) were visited once during the data collection period (from October 2013 to March 2014).

### 2.3 Data collection

The PBF impact evaluation used the HRITF survey instruments as a starting point and tailored them to the needs of the baseline survey and to the Burkinabe context [26]. Three different questionnaires were used in our study (Data in S1 Text).

The first one sought to assess health facilities and enabled, by direct and silent observation, to collect data on key aspects of their organization, their infrastructure and equipment, the availability of drugs, consumables and supplies (considering the pharmacy and/or drugstore located in the facility), their actual supervision (considering all the information included in the supervision book) etc. The second one (anonymous), dedicated to health workers, allowed investigating their role and responsibilities, their socio-demographic and professional characteristics, their satisfaction and motivation in relation to their work and their salaries (by using a scale score from 0 lowest to 10 highest) etc. The purpose of the last questionnaire was to gather information on how the consultations really took place, with the interviewer directly observing the interaction between patient and provider without being involved in. It was focused on IMCI protocols and a member of the survey team, therefore, directly observed the consultations, seeking to document which key processes of clinical care were carried out according to these guidelines (the provider was not aware of what kind of information that was collected). Data collection took place over five months, from October 15^th^, 2013 to March 15^th^, 2014.

Three health workers were interviewed in each health facility. For those with less than four health workers, all staff present at the facility was interviewed. We considered consultations of children aged between two months and 59 months and coming for visit in the PHC facilities. Four to five children (depending on the frequency of the consultations this day) presenting with a new condition (i.e. not for follow-up visits or routine) were observed during their consultation.

All data were collected by 30 investigators, accompanied by six direct supervisors (All were well trained during data collection trainings). A second level supervision was provided by three controllers and together, a team of supervisors from Centre MURAZ and University of Heidelberg provided a third level of control of the data collection. These different levels of quality control (collection, supervision and control) aimed at ensuring that similar results were obtained regardless of the investigator conducting the survey.

### 2.4 Study variables

The assessment of the adherence to guidelines (the dependent variables in our study) was based on a set of selected indicators, built and adapted (available on request as supplemental material) from the priority indicators for IMCI at health centres level proposed by WHO [3]. The potential factors influencing this adherence and by then, the quality -our independent variables - included PHC facilities and health workers characteristics. They were selected on the basis of a literature review and experts’ opinions including public health specialists and a paediatrician. The detailed composition of these indicators is described in Table 1. Algorithms and codifications were used to construct these variables when needed.

**Table 1:** Description of variables used

|  |  |
| --- | --- |
| **Variables used to assess quality of consultations according to IMCI protocols (dependent variables)** |  |
| ***Check (ask for and/or observe) for at least three general danger signs: (1)*** |  |
|  | • *Ability to drink or breastfeed* |
|  | • *Vomiting everything* |
|  | • *Convulsions* |
|  | • *Lethargic or unconscious* |
| ***Check and assess systematically for each of the five main signs/symptoms:*** |  |
|  | • *Cough or difficult breathing (2)* |
|  | • *Diarrhoea (3)* |
|  | • *Fever (4)* |
|  | • *Clinical anaemia (5)* |
|  | • *Ear problems (6)* |
| ***Check nutritional status (7)*** |  |
| *Check systematically Age* |  |
|  | • *Weight* |
|  | • *Height* |
|  | • *Swelling in the feet or ankles* |
| ***Check vaccination status (8)*** |  |
| ***Score of adherence (or IMCI Care score) = [summing of (1)+(2)+ (3)+(4)+ (5)+(6)+ (7)+(8), all having the same weight]*** |  |
| **Variables potentially associated with the adherence to IMCI care guidelines (Independent variables)** |  |
| ***Characteristics of the PHC facilities*** |  |
|  | • *Location (urban or rural)* |
|  | • *Availability of conveyance for referrals* |
|  | • *Number of supervision received* |
|  | • *Concomitant availability of all essential oral treatments [ORS, antibiotic against pneumonia (cotrimoxazole or amoxicillin), antibiotics against dysentery (cotrimoxazole or metronidazole), antimalarial (quinine tablet or ACT), vitamin A, iron, mebendazole and paracetamol]* |
|  | • *Concomitant availability of injectable drugs for pre-referral treatment [intramuscular antibiotic (ampicillin or ceftriaxone), quinine and benzyl penicillin]* |
|  | • Concomitant availability of equipment and supplies to provide comprehensive immunization service [*cold chain (cooler or ice-box or refrigerator or freezer), sterilizer and syringes (2 ml or 5 ml or 0.5 ml)*]. |
|  | • Availability of four vaccines (BCG, oral polio, DTP and measles) |
| ***Characteristics of health workers*** |  |
|  | Socio-demographic characteristics |
|  | • Age |
|  | • Sex |
|  | Socio-professional characteristics |
|  | • Qualification |
|  | • Seniority (number of years worked) |
|  | • IMCI training (period to which the nurses had the training) |
|  | • Salary satisfaction |

### 2.5 Data description and analysis

Data description and analysis were performed using statistical software STATA 12 and Excel 2007. The units of analysis were the consultations of children aged from 2 months to 59 months. Descriptive statistics for dependent as well as independent variables were depicted using common parameters, depending on the nature of the variable. Univariate and multivariate regressions analyses were used (after checking their adequacy to the models’ assumptions) to identify the predictors of adherence to IMCI guidelines. The association of each variable (dependent and independent) was checked (p<0.05 was used for significance). If there was not adequacy of the model assumption, the variable was left out. Only the factors with strength of evidence for an association are shown and discussed.

A logistic model (linear regression model) was used to assess the factors associated with each category of IMCI adherence: (i) check for at least three general danger signs, (ii) check and assess systematically each of the five main signs/symptoms, (iii) check nutritional status, (iv) check vaccination status. The adherence score (IMCI care score) represents the number of steps performed with success among the eight categories of the guidelines. A Poisson regression model was run to estimate the rate ratio of the associated factors.

## 3 Results

### 3.1 Characteristics of health facilities and health workers

The survey concerned 431 health facilities. 1,571 consultations of under five years old were observed. They were performed by 522 practitioners. Most of health facilities (92%) were located in rural areas and had no conveyances (considering the existence of available and functional ambulances) for referral (84%). During the last three months, health facilities were supervised on average three times by the hierarchy (either by the Health District and/or by the Regional Health Direction), but this number varied a lot from one facility to another, ranging from 0 to 13. Characteristics regarding drugs and vaccines availability are detailed in Table 2. One can notice that essential oral treatments, when considered individually (except for vitamin A), as well as injectable drugs and vaccines, were often available. But only one half of the health facilities had all the eight essential oral drugs. Equipment and supplies to provide a comprehensive immunization service were available all the time.

**Table 2:**
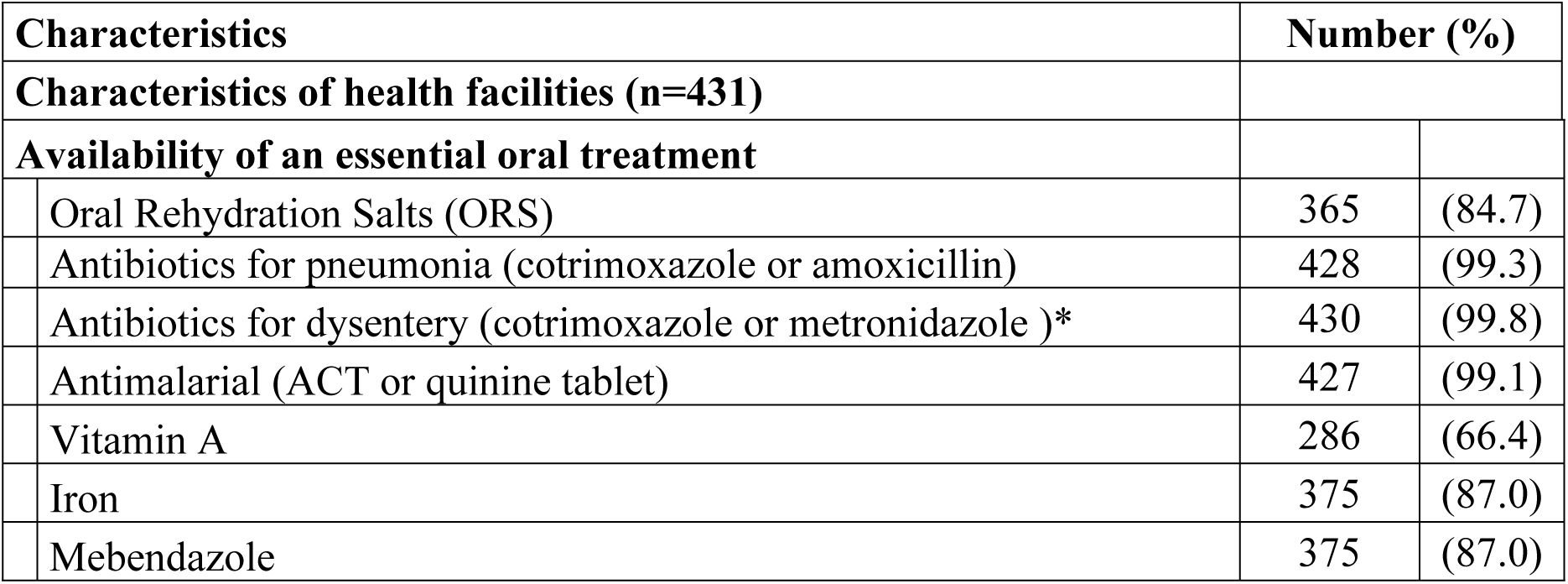
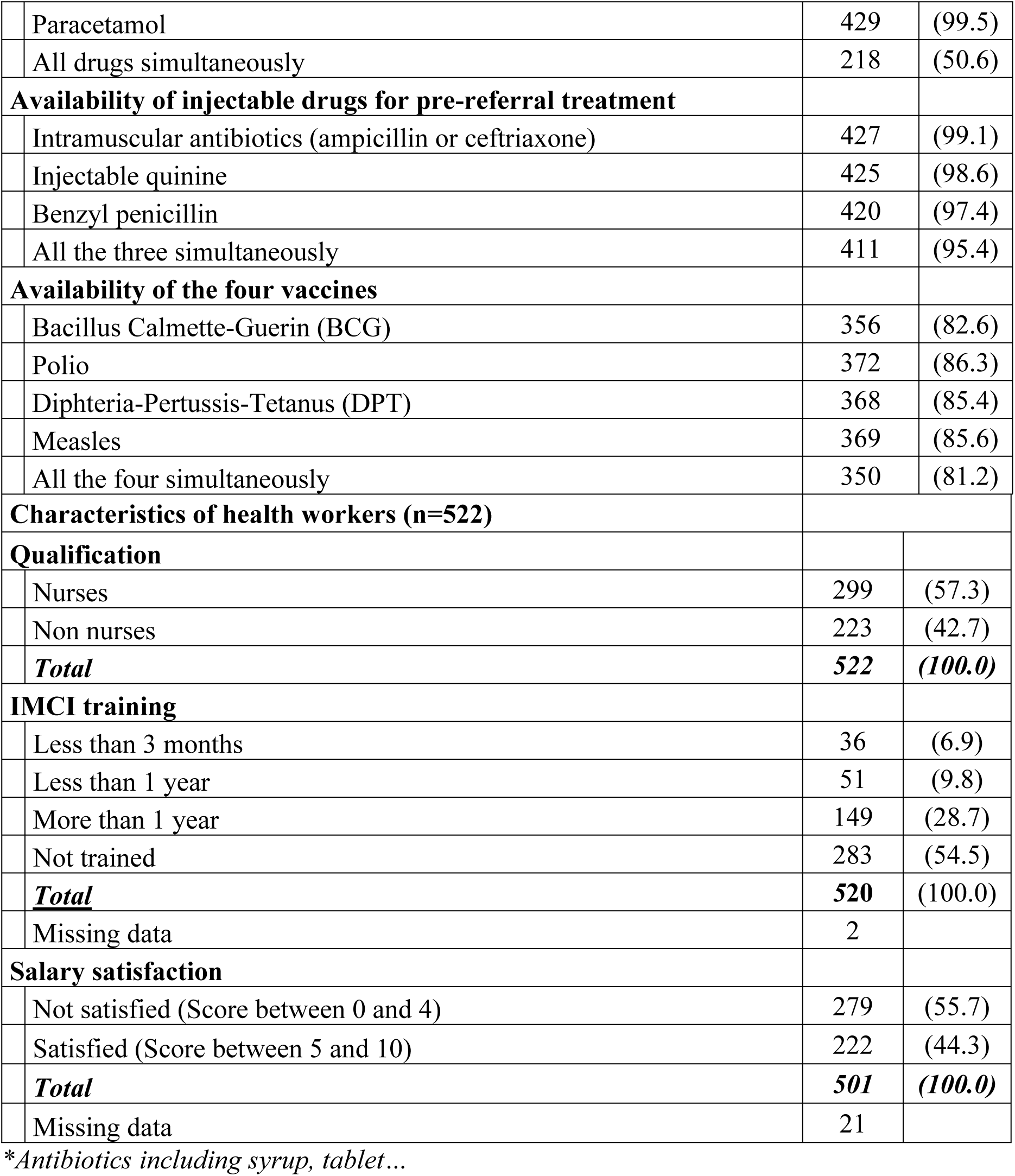
Characteristics of health facilities and health workers

| Characteristics |  | Number (%) |  |
| --- | --- | --- | --- |
| Characteristics of health facilities (n=431) |  |  |  |
| Availability of an essential oral treatment |  |  |  |
|  | Oral Rehydration Salts (ORS) | 365 | (84.7) |
|  | Antibiotics for pneumonia (cotrimoxazole or amoxicillin) | 428 | (99.3) |
|  | Antibiotics for dysentery (cotrimoxazole or metronidazole )* | 430 | (99.8) |
|  | Antimalarial (ACT or quinine tablet) | 427 | (99.1) |
|  | Vitamin A | 286 | (66.4) |
|  | Iron | 375 | (87.0) |
|  | Mebendazole | 375 | (87.0) |

|  |  |  |
| --- | --- | --- |
| Paracetamol | 429 | (99.5) |
| All drugs simultaneously | 218 | (50.6) |
| **Availability of injectable drugs for pre-referral treatment** |  |  |
| Intramuscular antibiotics (ampicillin or ceftriaxone) | 427 | (99.1) |
| Injectable quinine | 425 | (98.6) |
| Benzyl penicillin | 420 | (97.4) |
| All the three simultaneously | 411 | (95.4) |
| **Availability of the four vaccines** |  |  |
| Bacillus Calmette-Guerin (BCG) | 356 | (82.6) |
| Polio | 372 | (86.3) |
| Diphtheria-Pertussis-Tetanus (DPT) | 368 | (85.4) |
| Measles | 369 | (85.6) |
| All the four simultaneously | 350 | (81.2) |
| **Characteristics of health workers (n=522)** |  |  |
| **Qualification** |  |  |
| Nurses | 299 | (57.3) |
| Non nurses | 223 | (42.7) |
| **Total** | **522** | **(100.0)** |
| **IMCI training** |  |  |
| Less than 3 months | 36 | (6.9) |
| Less than 1 year | 51 | (9.8) |
| More than 1 year | 149 | (28.7) |
| Not trained | 283 | (54.5) |
| **Total** | **520** | **(100.0)** |
| Missing data | 2 |  |
| **Salary satisfaction** |  |  |
| Not satisfied (Score between 0 and 4) | 279 | (55.7) |
| Satisfied (Score between 5 and 10) | 222 | (44.3) |
| **Total** | **501** | **(100.0)** |
| Missing data | 21 |  |

\*Antibiotics including syrup, tablet...

The median age of the health workers was 33 (range 30 - 37) years. Almost the same number of health workers with less and more than five years of seniority performed the consultations (n=265 versus n=257). Children were more often examined by male health workers (68.4% of cases) but most of visits (57.1%) in urban areas were performed by the female ones. Details concerning health staff qualification, IMCI training status and degree of satisfaction regarding salary are given in Table in S2 Table. Up to 42.7% of consultations were carried out by staffs that were not qualified or trained to provide curative care to under-five (midwives/maieuticians, itinerant health workers – *agents itinerants de santé –* and assistant midwives – *accoucheuses auxiliaires –*) and among nurses, 34.9% were not trained to IMCI protocols, compared to 81.0% for other health professionals (non-nurses). More than half or the health workers (55.7%) were not satisfied with the salary they received.

### 3.2 Adherence to IMCI protocols

The detailed results of the assessment of this adherence are given in Table 3. But it is worth emphasizing that general danger signs were not investigated for more than half of observed consultations, except for the sign “vomiting everything”(51.4% of cases). Also, at least three general danger signs were systematically checked only for 13.9 % of consultations, while the five main symptoms/signs were all simultaneously checked and assessed in only 6.2% of cases. When considered individually, symptoms (except ears problems that were faintly examined) were checked for between 64.9% (if diarrhoea) and 83.8% (if fever) of cases. Assessment of their characteristics (e.g. duration, stridor for cough, blood in stools for diarrhoea etc.) varied greatly from one symptom/sign to another, but globally they were poorly performed. Indirect indicators (age, weight, height and oedema) were used to assess nutritional status and we can notice that the last two were the less investigated. Among the eight indicators we chose to give an adherence score to consultations (Table 1), three to five of them were achieved during 67.7% of consultations and none was performed during 2%.

**Table 3:** Adherence to IMCI guidelines

| **Components of consultations assessed (n=1,571)** |  | **Number** | **(%)** |
| --- | --- | --- | --- |
| **Check (Ask for and/or observe) for general danger signs** |  |  |  |
|  | Ability to drink or breastfeed | 620 | (39.5) |
|  | Vomiting everything | 807 | (51.4) |
|  | Convulsions | 262 | (16.7) |
|  | Lethargic or unconscious | 182 | (11.6) |
|  | At least three general danger signs checked | 219 | (13.9) |
| **Cough** |  |  |  |
|  | Actually asked for | 1175 | (74.8) |
| **Cough assessment (Cough present in 604 cases)** |  |  |  |
|  | Ask for duration | 545 | (90.2) |
|  | Check for stridor | 143 | (23.7) |
|  | Respiratory rate reckoned | 335 | (55.5) |
|  | Auscultation performed | 333 | (55.1) |
|  | All these four elements simultaneously checked | 89 | (14.7) |
| **Diarrhoea** |  |  |  |
|  | Actually asked for | 1020 | (64.9) |
| **Diarrhoea assessment (Diarrhoea present in 368 cases)** |  |  |  |
|  | Ask for duration | 350 | (95.1) |
|  | Ask for blood in stools | 269 | (73.1) |
|  | Skin pinch to check for dehydration | 200 | (54.3) |
|  | All these three elements simultaneously checked | 154 | (41.8) |
| **Fever** |  |  |  |
|  | Actually checked (Asked for and/or measured temperature) for | 1316 | (83.8) |
| **Search for the cause of fever (Fever present in 644 cases)** |  |  |  |
|  | Ask for duration | 569 | (88.3) |
|  | Ask for history of measles | 35 | (5.4) |
|  | Eyes examined | 517 | (80.3) |
|  | Nose examined | 147 | (22.8) |
|  | Throat examined | 92 | (14.3) |
|  | Check for skin rash | 106 | (16.5) |
|  | All these six elements simultaneously checked | 9 | (1.4) |
|  | Fontanel examination for children under 8 months of age (394 patients) | 25 | (6.3) |
| **Ears' problems** |  |  |  |
|  | Inside examination | 245 | (15.6) |
|  | Outside examination | 284 | (18.1) |
|  | Ask for pain or observe discharge | 385 | (24.5) |
|  | Complete inspection (comprising the above three elements) | 178 | (11.3) |
| **Check for clinical anaemia (look at eyes, palms or soles)** |  | 1104 | (70.3) |
| **Five main signs/symptoms (cough, diarrhoea, fever, ears problems and clinical anaemia) checked for and assessed** |  | 98 | (6.2) |
| **Check for vaccination status (look at the vaccination card)** |  | 665 | (42.3) |
| **Nutritional assessment** |  |  |  |
|  | Asked for age | 1440 | (91.7) |
|  | Weight measured | 1062 | (67.6) |
|  | Height measured | 374 | (23.8) |
|  | Check for oedema | 720 | (45.8) |

### 3.3 Factors associated with adherence to IMCI

A comprehensive summary of multivariate analysis results shows that factors negatively associated with adherence according to IMCI guidelines were: (i) other qualification than nurses, (ii) female practitioners, (iii) non-satisfaction with salary. The factor positively associated was IMCI training. The details of results for each dependent variable, for univariate as well multivariate analysis, are shown in Table 4.

**Table 4:** Factors associated with adherence to IMCI guidelines

| Variables | Crude RR |  |  | Adjusted RR |  |  |
| --- | --- | --- | --- | --- | --- | --- |
|  | RR | 95% CI | p | RR | 95% CI | p |
| **General danger signs** |  |  |  |  |  |  |
| **Sex** |  |  |  |  |  |  |
| Male | 1 |  |  | 1 |  |  |
| Female | 0.67 | 0.48-0.93 | 0.017 | 0.74 | 0.53-1.04 | 0.081 |
| **Practitioner** |  |  |  |  |  |  |
| Nurse | 1 |  |  | 1 |  |  |
| Non nurse* | 0.61 | 0.45-0.83 | 0.001 | 0.72 | 0.50-1.02 | 0.065 |
| **IMCI training** |  |  |  |  |  |  |
| No training | 1 |  |  | 1 |  |  |

| Variables |  | Crude RR |  |  | Adjusted RR |  |  |
| --- | --- | --- | --- | --- | --- | --- | --- |
|  |  | RR | 95% CI | p | RR | 95% CI | p |
|  | Trained | 1.50 | 1.13-2.00 | 0.005 | 1.24 | 0.89-1.73 | 0.197 |
| **Signs and symptoms** |  |  |  |  |  |  |  |
| *Availability of a mean of conveyance* |  |  |  |  |  |  |  |
|  | No | 1 |  |  | 1 |  |  |
|  | Yes | 1.77 | 1.10-2.86 | 0.020 | 1.58 | 0.98-2.55 | 0.063 |
| *Availability of injectable drugs* |  |  |  |  |  |  |  |
|  | No | 1 |  |  | 1 |  |  |
|  | Yes | 0.41 | 0.20-0.85 | 0.017 | 0.36 | 0.16-0.77 | 0.009 |
| **IMCI Training** |  |  |  |  |  |  |  |
|  | No training | 1 |  |  | 1 |  |  |
|  | Trained | 2.61 | 1.69-4.03 | <0.001 | 2.61 | 1.71-3.98 | <0.001 |
| **Nutritional assessment** |  |  |  |  |  |  |  |
| *Practitioner* |  |  |  |  |  |  |  |
|  | Nurse | 1 |  |  | 1 |  |  |
|  | Non nurse | 0.68 | 0.49-0.93 | 0.016 | 0.66 | 0.48-0.93 | 0.010 |
| **Location** |  |  |  |  |  |  |  |
|  | Urban | 1 |  |  | 1 |  |  |
|  | Rural | 0.61 | 0.37-0.99 | 0.046 | 0.62 | 0.38-1.01 | 0.056 |
| *Availability of an essential oral treatment* |  |  |  |  |  |  |  |
|  | No | 1 |  |  | 1 |  |  |
|  | Yes | 2.14 | 1.55-2.94 | <0.000 | 2.17 | 1.58-3.00 | <0.001 |
| **Vaccination status** |  |  |  |  |  |  |  |
| *Supervision* |  |  |  |  |  |  |  |
|  | 3 or more in the last three months | 1 |  |  | 1 |  |  |
|  | Less than 3 in the last three months | 0.78 | 0.63-0.97 | 0.023 | 0.79 | 0.64-0.98 | 0.034 |
| *Availability of an oral essential treatment* |  |  |  |  |  |  |  |
|  | No | 1 |  |  | 1 |  |  |
|  | Yes | 1.33 | 1.09-1.63 | 0.004 | 1.30 | 1.06-1.58 | 0.010 |
| **IMCI care score\*\*** |  |  |  |  |  |  |  |
| **Sex** |  |  |  |  |  |  |  |
|  | Male | 1 |  |  | 1 |  |  |
|  | Female | 0.89 | 0.85-0.94 | <0.001 | 0.91 | 0.86-0.95 | <0.001 |
| **Practitioner** |  |  |  |  |  |  |  |
|  | Nurse | 1 |  |  | 1 |  |  |
|  | Non nurse | 0.88 | 0.84-0.92 | <0.001 | 0.93 | 0.88-0.97 | 0.002 |
| **IMCI Training** |  |  |  |  |  |  |  |
|  | No training | 1 |  |  | 1 |  |  |
|  |  | RR | 95% CI | p | RR | 95% CI | p |
|  | Trained | 1.12 | 1.07-1.16 | <0.001 | 1.06 | 1.01-1.11 | 0.024 |
| *Satisfied by salary* |  |  |  |  |  |  |  |
|  | Satisfied | 1 |  |  | 1 |  |  |
|  | Not satisfied | 0.95 | 0.91-0.99 | 0.013 | 0.95 | 0.91-0.99 | 0.022 |

\* Non nurse includes midwives/maieuticians, itinerant health workers – *agents itinérants de santé* – and assistant midwives – *accoucheuses auxiliaires*
\*\* Results derived from a Poisson regression model. Data are Rate Ratio with 95% CI and p-value.

## Discussion

This study presents factors associated with the adherence to IMCI guidelines (with extrapolation to quality of care) in Burkina Faso PHC facilities. The focus was on the clinical and health system strengthening components of IMCI, highlighting challenges encountered by Burkina Faso like many other sub-Saharan countries when implementing the strategy [27,28].

In Burkina Faso, consultations are often carried out by staff other that nurses such as midwives, itinerant health workers (AIS)… Shortage of skilled health personnel is an important issue and is a recurrent problem in most of the LMICs [29,30]. This often leads to an overwhelming workload for nurses and to give them a hand, “unqualified” or trained for other tasks staffs – such as assistant midwives, itinerant health workers – usually provide curative care to under-five children. In fact, the main role of assistant midwives in Burkina Faso context (we consider midwives roles being obvious) is to: (i) practice eutocic deliveries as well as pre and post-natal consultations, (ii) refer risky pregnancies as well as complicated and obstructed labours, (iii) contribute to health education, specifically for family planning, (iv) manage and supervise village midwives. Itinerant health workers, meanwhile, have to: (i) advice and assist patients and the community regarding hygiene issues, (ii) conduct home visits to identify sick people, pregnant women, new-borns and infants, in order to refer them to a health facility, (iii) mobilize the community for preventive activities, (iv) manage and supervise community health workers [31]. Regarding IMCI training, many studies already reported its importance, showing that those trained performed better than those who were not [5,6,17,32,33]. So, building skills is crucial in IMCI strategy, both in terms of initial and continuing trainings.

Our results showed that staff other than nurses were less trained to IMCI despite that according to WHO, IMCI could be performed by doctors, nurses and other health workers who give care to sick children [34]. So, IMCI training should target any health worker in Burkina Faso (Even though, the health policy gives priority for training to the nurses) given the context of shortages in human resources for health leading “unqualified” staff to provide curative care to children. The main constraints remain resource availability as training processes are expensive and sometimes considered cumbersome by stakeholders [29]. One solution could be to introduce IMCI training in the curricula of health training schools, which would reduce costs (costs effectiveness), improve basic skills of health workers, and quickly provide health facilities with competent staff with a common understanding. Examples are provided from ten countries (Cambodia, China, Fiji, Kiribati, the Lao People’s Democratic Republic, Mongolia, Papua New Guinea, the Philippines, Solomon Islands and Viet Nam) where pre-service IMCI education was successfully implemented, varying from one country to another, in the medical and nursing schools [35]. In Ethiopia (where pre-service IMCI education had also been implemented), the most preferred teaching style was the mixed approach including group discussion and demonstration [36]. There is an opportunity to learn from these countries and implement similar approach (with local adaptation) in Burkina Faso. The health system would in this case only have to organize refreshment training and follow up.

We can also notice that the working conditions/environment is not always supportive in terms of incentives for many health workers. Indeed, most of them were not satisfied with their wages. Many studies already evidenced the crucial role of extrinsic motivation regarding provision of quality health care in low-resource settings [19,37]. Moreover, poor remunerations could undermine intrinsic motivation because, among others, decent pay is seen as an important way of giving recognition to health workers and would enable them to face living expenses [19]. In particular in Burkina Faso as it is acknowledged that wages are lower compared to other sub-Saharan Africa countries without great differences in living standards [37].

Lack of equipment and supplies in health facilities, as well as poor organization, could also demoralize staff. In our study, drugs were usually available as well as vaccines and equipment/supplies to provide a comprehensive immunization service. But it is worth noting that most of health facilities lacked conveyances and big differences existed among health facilities regarding the supervision they benefitted from their hierarchy, the numbers are ranging from zero to 13 in the last three months. Beyond an issue of health system organization, these disparities could be explained by a difficult geographical access (some being landlocked during rainy seasons).

Supervision also requires mobilizing enormous resources, particularly human, financial and logistical ones which are not always available. So, specific supervisions dedicated to IMCI are not achieved and integrated supervisions are preferred even if they are less effective (mainly because of the multiplicity of items that must be taken into account), as widely acknowledged by many studies [29,30]. This is compounded by the fact that many health facilities were located in rural (and probably remote areas which reflects the distribution of health facilities in the country). Yet, regular monitoring and supervision of health providers are indispensable if we want efficient health systems, especially as there is task shifting for staff working at peripheral level.

Regarding the adherence to IMCI guidelines and the factors associated with, the general danger signs were not always checked; at least three of these signs being sought in only 13.9% of consultations. Any child with such a sign needs urgent care and hospitalization and should be swiftly referred after a “pre-transfer” treatment. Main symptoms or signs, if they were checked for, were not always assessed while we know that this is indispensable to perform all clinical assessments in order to classify them for providing the most accurate treatment [3,34,38]. Moreover, vaccination and nutritional status were not always well investigated.

Many factors could explain these results and suggest that the care provided are of poor quality or at least not optimal. In addition to their limited knowledge (skills issues), we can raise (from our clinical observations during the study) negligence issues, some health workers are more relying on their own experience and capacities instead of following IMCI guidelines rigorously. Workload issues could also be raised, not necessarily related to a high number of patients, but more to the experience of being more or less constantly on duty, thereby leading to too little time to rest and hence, a physical overload [19] and also the time needed to fill in the registers for the health information system.

Finally, we found enablers and barriers factors pertaining to health facilities and health workers characteristics which were linked to the adherence of IMCI and therefore, to quality of child health care, sometimes counter-intuitively. This is the case for the availability of injectable drugs, the supervision of health facilities and being a female health worker, which were negatively associated, the first with the main symptoms/signs, the second with the check of vaccination status and the last with the consultation adherence score. Constant availability of injectable drugs could entail misuse by health workers; if they are not motivated to work, they could use them and refer children without taking time to thoroughly examine them, as our results strongly suggest it. For supervision, we presume that they were not efficient as they usually do not focus on IMCI. Results for female workers need more investigation, especially a qualitative study would be needed to understand better the reasons behind.

Lange *et al* [19], while trying to understand why clinicians do not adhere more consistently to IMCI guidelines in Tanzania, found two main reasons: a lack of capacity or a lack of motivation. These two explanations seem also consistent with our findings, beyond contextual specificities. Definitely, the description of PHC facilities and health workers’ characteristics suggests some limited knowledge of the strategy (and thus, an actual need of capacity building) and a demotivating working environment and/or conditions.

These last authors (Lange et al) described that the direct observation technique, beyond its subjectivity, may induce health workers to improve their performance above normal levels, thus introducing bias in data collection (a Hawthorne effect).

A limitation of our study is that we did not assess treatment quality and outcomes such morbidity (quality of care through adherence is actually a complex issue), even if we made the hypothesis that a positive correlation exists between a good consultation and a treatment quality, and so a good health outcome, as Donabedian’s framework of structures – processes – outcomes [39] suggests. But this is not always granted [40]. Indeed, “good results” can occur after inadequate treatment (e.g. a fever due only to malaria but treated concomitantly with anti-malarial drugs and a broad spectrum antibiotic, resulting in a recovery of the patient). Conversely, poor outcomes (patient deaths) are consistent with achievement of excellent care processes in high-quality care structures (e.g. treatment of incurable cancer). Another limitation was the fact that we used secondary data, as this may prevent collecting relevant variables that would more inform our results. Moreover, direct observations could introduce bias as health workers knowing that they are observed, would be prone to change their usual habits and try to perform better (Hawthorne effect). Finally, even if we conducted our study in six regions on 13, all health districts were not represented. That might not represent all IMCI adherence and quality of child health care in the country.

## 5 Conclusions

The IMCI strategy is a powerful tool that could help reduce under-five mortality and morbidity in developing countries, especially in Burkina Faso where indicators are still lagging despite some improvements in the last years. Our study highlights that there is still considerable room for improvement in its clinical component. Adherence to IMCI guidelines was found poor in the study, driving to a poor quality of care. Indeed, while some symptoms such as cough, diarrhoea or fever have been relatively well checked for, their assessment that would enable classify the case in order to give the appropriate care to children was not often realized. In addition, the general danger signs were not checked for in over half the consultations while their presence indicates an emergency and a need of referral. The systematic search for all elements as recommended in the protocol was rarely done. Factors such as, other qualification than nurses, female practitioners, non-satisfaction with salary, were negatively associated with IMCI care while IMCI training was positively.

If further inquiries are needed to better understand their actual influence, a prospective solution would come from health workers’ capacity building and the improvement of their extrinsic and intrinsic motivation. But this should ideally be part of a comprehensive approach aiming at strengthening health system as a whole. A new strategy, based on information technology is now under assessment in Burkina Faso [41]. This could, like the PBF approach, improve adherence to IMCI guidelines in the PHC facilities.

## Acknowledgments

We would like to thank all the Burkina Faso RBF impact evaluation team, the Ministry of Health of Burkina Faso, the World Bank, the Heidelberg University, the Montreal University and the Centre MURAZ. We would also like to thank the Centre MURAZ data management team for their support.

## Supporting Information Captions

**S1 Table 1.** Description of variables used

**S2 Table 2.** Characteristics of health facilities and health workers

**S3 Table 3.** Adherence to IMCI guidelines

**S4 Table 4.** Factors associated with adherence to IMCI guidelines

**S1 Data.** Data from the Baseline study – Impact Evaluation of the Performance Based Financing - available at http://microdata.worldbank.org/index.php/catalog/2761

**Authors’ contributions**
Regarding contribution to authorship, SAT participated in the conceptualization, methodology, data curation, formal analysis, and writing of the manuscript. SMAS participated in the methodology, investigation, software, data curation, formal analysis, and the writing of the manuscript. JAK participated in the conceptualization, methodology, supervision and the writing of the manuscript. JLK contributed in the investigation, and reviewed the manuscript. PJR contributed in the investigation, the funding acquisition and reviewed the manuscript. HH was involved in the investigation, the funding acquisition, project administration, and in the review of the manuscript. NM conceptualized the study, contributed in the funding acquisition, reviewed and edited the manuscript.
All authors read and approved the final manuscript.

